# Telomere interactions and structural variants in ALT cells revealed with TelSPRITE

**DOI:** 10.1101/2024.11.22.624895

**Authors:** David G. Wilson, Sarah F. Clatterbuck Soper, Marbin A. Pineda, Robert L. Walker, Paul S. Meltzer

## Abstract

Acquisition of a telomere maintenance mechanism is essential for cancer cells. In a minority of tumors, telomeres are lengthened via Alternative Maintenance of Telomeres (ALT), a telomerase-independent pathway based on homologous recombination. ALT tumors have heavily rearranged genomes with many structural variants containing telomere repeats. To better understand the genetic evolution of these tumors, we seek to determine if certain genomic loci tend to spatially associate with telomeres in ALT and are especially liable to experience telomere recombination events as a result. Assays that reveal close spatial associations between genomic loci, such as SPRITE and Hi-C, have enabled extensive exploration of genomic spatial organization. However, as analysis pipelines for these next-generation sequencing-based assays typically discard reads aligning to repetitive elements, little is known about the spatial arrangement of telomeres and other repetitive loci in the nucleus. Here, we present TelSPRITE, a novel approach to extracting telomere contact frequencies from SPRITE data. We identify reads containing telomere repeats and sort them into a single bin, quantifying spatial contacts between the telomere bin and the rest of the genome. Our analysis reveals a strong dependency of telomere contact frequency on chromosomal distance from the telomere, consistent with the known effect of linear distance on 3-dimensional spatial contacts. Telomere contacts are also strongly enriched near centromeres, a phenomenon that may be reflective of spatial clustering of heterochromatic regions. ALT cell lines are globally enriched for telomere content and display distinctive intrachromosomal spikes in telomere contact frequency. Our customized analysis of long read sequencing data suggests that loci with high telomere contact frequencies represent structural variants containing telomere repeats in ALT cells. Collectively, our results demonstrate general principles of telomeric spatial organization while also profiling the spectrum of genomic rearrangements in ALT cells.

## Introduction

Cancer cells must acquire a telomere maintenance mechanism to counteract the shortening of telomeres that occurs with successive cell divisions.^1^ While most tumors express telomerase to achieve telomere lengthening, in about 10-15% of tumors telomeres are instead lengthened via a homologous recombination-based pathway called Alternative Lengthening of Telomeres (ALT).^2–4^ ALT is strongly associated with defects in the ATRX/DAXX histone chaperone complex, which deposits histone variant H3.3 at telomeres.^5–7^ ATRX/DAXX plays an important role in unwinding telomeric G-quadruplexes, and loss of ATRX/DAXX function is associated with replication stress and DNA damage at telomeres.^8,9^ In ALT, dysfunctional telomeres aggregate at PML bodies, where they are lengthened via break-induced replication.^10–12^

ALT tumors have heavily rearranged genomes and many structural variants containing telomere sequence.^13^ Given that ALT telomeres actively undergo homologous recombination and that telomere sequence is found at ectopic sites in ALT tumors, we hypothesized that recombination events between telomeres and other genomic loci drive genomic rearrangements in ALT cells. However, little is known about what regions of the genome are particularly likely to undergo these recombination events. We hypothesized that spatial proximity may be a factor: Genomic loci that are spatially engaged with telomeres may be liable to recombine with them. To explore this hypothesis, we examined telomere spatial arrangement with a particular focus on ALT cells.

Several methods for determining spatial associations between genomic loci currently exist and have different sets of advantages. These methods include Hi-C, SPRITE, and GAM.^14–16^ GAM infers genomic interactions based on which loci are most likely to colocalize in nuclear cryosections,^16^ whereas Hi-C and SPRITE identify genomic interactions by determining which genomic loci are most likely to crosslink to each other. Hi-C follows chromatin crosslinking and fragmentation with a proximity ligation step which attaches crosslinked genome fragments end-to-end before sequencing, while SPRITE utilizes an iterative barcoding procedure to identify clusters of crosslinked DNA fragments. SPRITE has the advantage of enumerating complex, multiway interactions that occur in crosslinked clusters containing many genome fragments, whereas Hi-C is mostly limited to pairwise interactions given the short read lengths produced by next-generation sequencing.^14,15^

Despite significant progress in elucidating the spatial arrangement of much of the genome, understanding of the arrangement of repetitive DNA regions in the nucleus, such as telomeres, remains limited. Analysis pipelines for 3-dimensional genomics assays typically mask repetitive genomic loci since next-generation sequencing reads do not align to them uniquely. However, while establishing complete 2-dimensional contact frequency maps of these loci is not possible with current methods, 1-dimensional profiling of the contact frequency of all DNA fragments consisting of a given repetitive sequence against the rest of the (non-repetitive) genome should still be possible with modifications to the data analysis pipelines.

To reveal genomic interactions with telomeres, we developed TelSPRITE, a computational framework that quantifies the frequency of spatial contacts with telomeres genome-wide. We altered the existing SPRITE pipeline by adding a step that identifies sequenced reads containing canonical and variant telomere repeats.^17^ Telomeric reads were then assigned to an artificial genomic locus. The telomere locus was included during generation of the SPRITE contact frequency matrix to determine the contact frequency between telomeres and other genomic loci.

Our analysis of telomere spatial distribution across multiple cell types reveals general principles that govern telomere contacts. We uncover a strong dependance of telomere contact frequency on chromosomal distance from the telomere, a phenomenon consistent with the finding that linearly proximal genomic loci display increased contacts in 3-dimensional space in SPRITE and Hi-C.^14,15^ Furthermore, frequent and substantial spikes in telomere contact frequency are seen near centromeres, an observation consistent with spatial clustering of heterochromatic regions. A broader relationship between telomere contacts and heterochromatin is revealed in at least some cell lines.

In ALT cells, we observe an overriding dominance of ectopic telomere repeats over the telomere contact pattern. ALT cell lines share many basic principles of telomere spatial organization with non-ALT cell lines, including linear dependance of contacts and enriched interactions with centromeres, but are globally enriched for telomere content, have frequent aberrant spikes in telomere contact frequency internal to chromosomes explained by structural variants containing telomere repeats, and exhibit less clear evidence of enhanced telomere contacts with non-centromeric heterochromatin. Furthermore, rapid suppression of ALT globally reduces telomere content, which may imply that SPRITE captures contacts of ALT-specific extrachromosomal telomere DNA species.

Our results show that key aspects of telomeric spatial organization are consistent regardless of ALT status even as TelSPRITE reveals an extensive pattern of ectopic telomere repeats and global telomere content enrichment to be characteristics that distinguish ALT-positive cell lines. Furthermore, we provide a useful and easily adaptable computational framework for extracting a contact pattern for a repetitive genomic element from 3-dimensional genomics data, which may aid in future studies seeking to examine spatial contacts of repetitive regions.

## Results

### SPRITE successfully captures known principles of genome organization

To interrogate the spatial organization of telomeres in ALT versus non-ALT cells, we performed SPRITE on two ALT-positive osteosarcoma cell lines (U2OS and G292-iDAXX) and one ALT-negative osteosarcoma cell line (HOS). G292 has a chromosomal translocation that causes loss of DAXX function; we previously developed G292-iDAXX to enable rapid suppression of the ALT mechanism via inducible expression of wild-type DAXX.^18^ We include G292-iDAXX with five days of ALT suppression to determine if rapid ALT suppression rearranges telomere contacts. We also include our reanalysis of the published SPRITE dataset from the lymphoblastoid cell line GM12878^15^ using the TelSPRITE pipeline to extract telomere contacts.

The SPRITE protocol was performed as described without modification.^17^ Our datasets encompass a wide range of cluster sizes (Figure S1A). Similar to the published GM12878 dataset, we observe known patterns of genomic spatial arrangement, including a dependance of contacts on linear distance (Figure S1B) and a prominent “plaid” pattern in Pearson correlation matrices of intrachromosomal contacts characteristic of A/B chromatin compartmentalization (Figure S1C).^14^

### Modifications to the SPRITE computational pipeline enable determination of telomere contact frequencies

Changes to the SPRITE data analysis pipeline were needed to extract telomere contacts (Figure 1A). Briefly, we added a step prior to read alignment that divided telomeric and non-telomeric reads into separate fastq files. Telomere reads were then assigned to an artificial genomic locus (called “chrT”) whereas the non-telomeric reads were aligned to the hg38 reference genome. The two sets of reads were then recombined before generating a cluster file containing groups of reads with shared barcodes. The 2-dimensional contact frequency matrix was then generated with a row corresponding to the locus chrT; the contact frequencies in this row could be easily extracted. For matrix generation, a binning resolution of 1 Mb was used, meaning that our telomere contact frequencies are calculated over successive 1 Mb sections of each chromosome.

**Figure 1.**
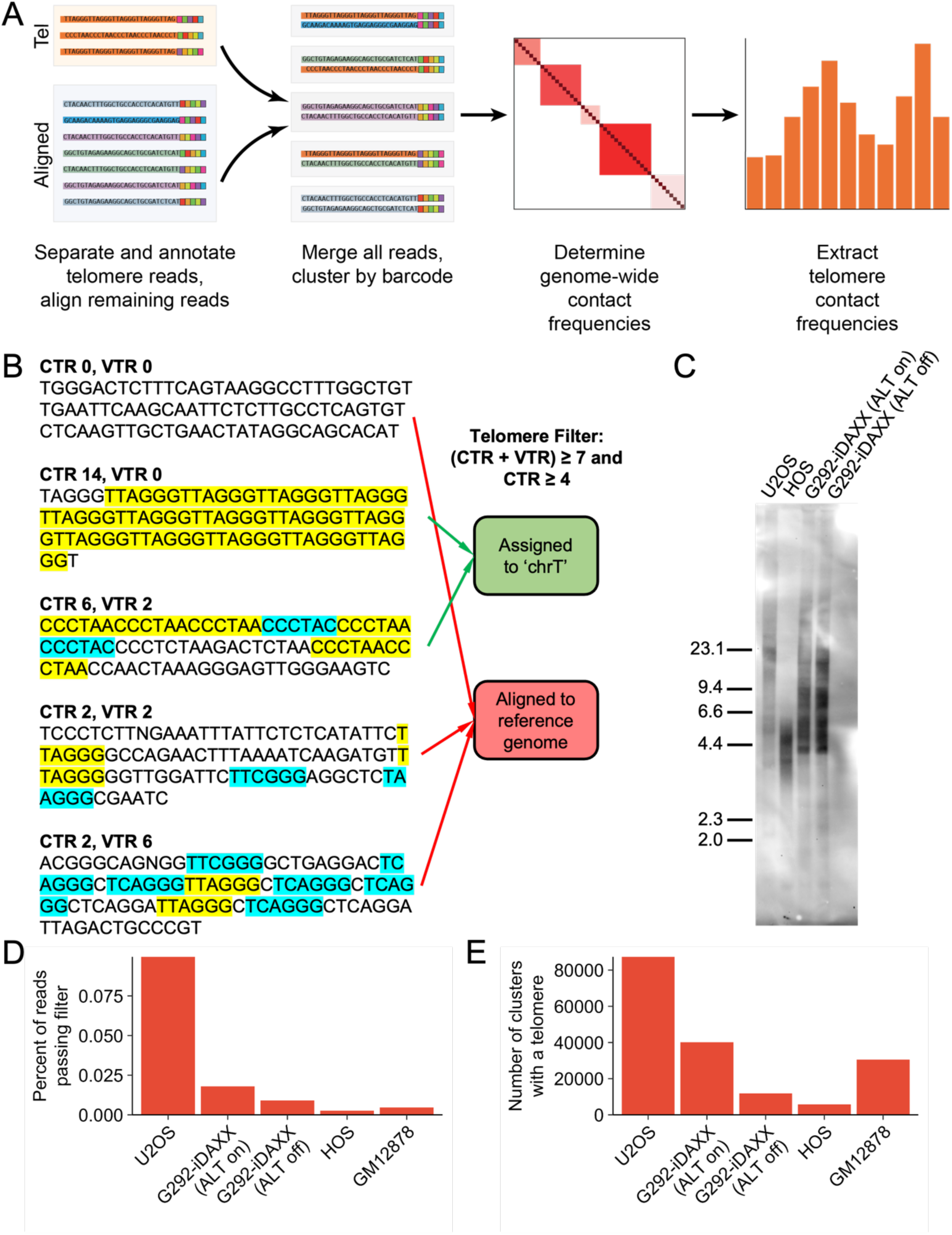
Determination of telomere contact frequencies with SPRITE. **(A)** Procedure for quantifying telomere contact frequencies using SPRITE. **(B)** Process for identifying telomere reads. The numbers of canonical (yellow) and variant (blue) repeats in a SPRITE read were counted, and if the specified thresholds for total repeats and canonical repeats were both satisfied, the read was assigned to the artifical locus chrT. Reads that failed either or both thresholds were aligned against the reference genome as normal. **(C)** ALT-positive U2OS and G292-iDAXX exhibited longer and more heterogenous telomere lengths on a telomere Southern blot than ALT-negative HOS. Rapid ALT suppression in G292-iDAXX did not cause measurable telomere length attrition. **(D)** Telomere content by cell line. Bar plot displays the percentage of reads that passed the filter described in (B) in each cell line. **(E)** Number of SPRITE clusters containing at least one telomere in each data set. The quantity of telomere-containing clusters depends on multiple factors, including the telomere content of the cell line and the number of unique SPRITE clusters that were produced and sequenced.

Telomere filtering was initially attempted at three different stringency thresholds. To count as a telomere, reads were required to contain 4, 7, or 14 telomere repeats for “low,” “medium,” and “high” stringency respectively. Sequencing of SPRITE reads yields about 85-90 base pairs of genomic DNA to align, so the maximum possible number of telomere repeats in a read will be around 14, though some fragments are smaller. Human telomeres, particularly in ALT-positive cell lines, are also known to feature variant telomere repeats that differ slightly from the canonical sequence (TTAGGG/CCCTAA).^19,20^ Therefore, we were permissive towards some variant repeats, but still required minimum numbers of canonical repeats (2, 4, and 7 for low, medium, and high stringencies respectively). A list of variant repeats is provided in Table S1.^21^

The impact of filtering conditions on the contact pattern was minimal (Figure S2). For following analyses, the “medium” stringency conditions were selected. SPRITE reads that satisfied both filtering conditions (7 or more total repeats, including both canonical and variant repeats, and 4 or more canonical repeats) were considered to contain telomeric sequence and assigned to the chrT locus; reads that failed either or both conditions were assumed to not originate from telomeres and were aligned to the hg38 reference genome to determine their genomic location (Figure 1B).

U2OS and G292-iDAXX exhibit substantially longer telomeres than HOS in agreement with the fact that ALT cells are known to generally have telomeres with long and heterogenous lengths (Figure 1C).^2^ Consistent with the difference in telomere lengths, a substantially higher percentage of reads pass the telomere filter in the ALT-positive cell lines (Figure 1D). The number of SPRITE clusters containing at least one read that meets the medium stringency filtering criteria is in the thousands or tens of thousands for all datasets (Figure 1E).

### Telomere contact frequency is high near chromosome ends

Spatial contacts in Hi-C and SPRITE are most likely between linearly proximal loci.^14,15^ Consistent with a correlation between chromosomal distance and high likelihood of 3-dimensional contacts, telomere contact frequencies are often very high at the chromosome end near the telomere (Figure 2A). Averaging the telomere contacts of the set of all SPRITE bins that share a given distance from the chromosome end shows a precipitous drop in telomere contact frequency with increasing chromosomal distance from the telomere (Figure 2B). Examining individual SPRITE clusters for telomere contacts at high resolution demonstrates that contacts disproportionately occur near the very end of the chromosome arm (Figure 2C).

**Figure 2.**
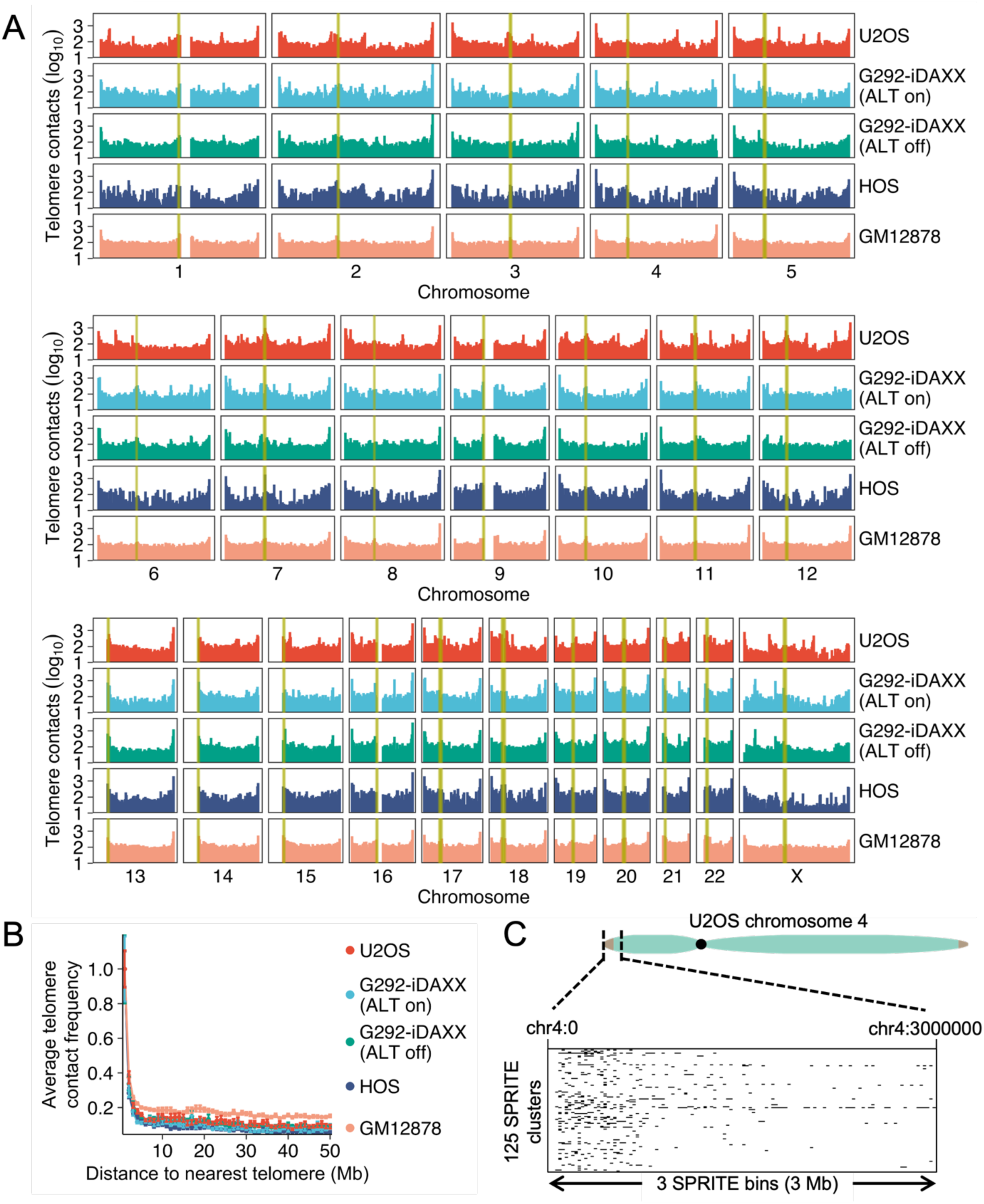
Telomere contact frequencies are heavily driven by chromosomal distance. **(A)** Genome-wide telomere contact pattern for U2OS, G292-iDAXX with active ALT, G292-iDAXX with suppressed ALT (suppressed via five days of induced overexpression of WT DAXX), HOS, and GM12878. Yellow highlighting denotes SPRITE bins containing centromere sequence. **(B)** Telomere contact signal decreases with increasing distance from the telomere. Telomere contact frequency for each cell line was averaged across each chromosome arm at a given distance from the nearest telomere (based on the hg38 reference genome) and expressed as a ratio to the average contact frequency at a distance of 1 Mb. Bins with a contact frequency of exactly zero were assumed to lack mappable DNA and excluded from the calculation. Error bars represent standard error. **(C)** The U2OS chromosome 4 p-arm provides an example of disproportionate telomere contacts near the chromosome end. Each row of the cluster plot represents a telomere-containing SPRITE cluster that has a size of 100 or less and at least one contact in the first 3 Mb of chromosome 4. Black lines show loci contacting telomeres in these clusters at a resolution of 25 kb.

For analyses examining telomere contacts with centromeres, other heterochromatic loci, and interstitial telomere sites, we “distance-normalized” the telomere contact frequency in each SPRITE bin by dividing the telomere contact frequency by the average contact frequency for bins at the same distance from the nearest telomere. This manipulation allowed us to better examine other factors impacting telomere contact frequency besides the distance from the chromosome end, which would otherwise be the dominant signal.

### Telomeres and centromeres spatially associate

A unifying feature of telomere contact frequencies in all cell lines examined was frequent enrichment near centromeres (Figure 2A). For example, U2OS exhibited a large spike in telomere contacts around the centromere of chromosome 7 (Figure 3A). To assess this trend genome-wide in all cell lines, we determined which SPRITE bins correspond to chromosomal segments with centromere sequence based on the hg38 centromere sequence annotation track on the UCSC Genome Browser.^22,23^ SPRITE bins overlapping centromere sequence exhibited substantially larger distance-normalized telomere contact frequencies than those that did not (Figure 3B).

**Figure 3.**
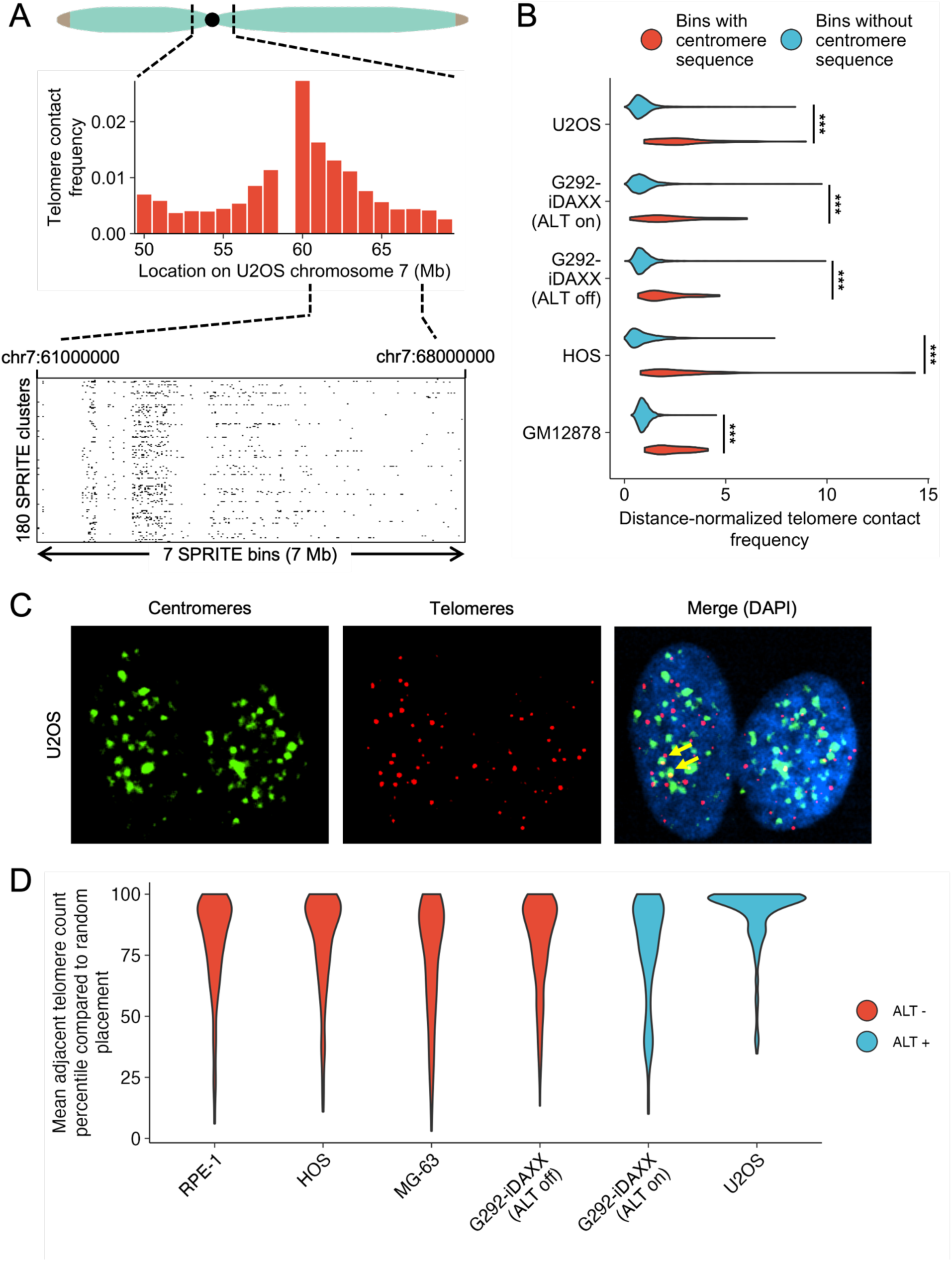
Telomeres and centromeres display a spatial association. **(A)** U2OS chromosome 7 exhibits a high telomere contact frequency near the centromere. Each row of the plot shows an individual SPRITE cluster with a size of 100 or less that has a telomere and another contact in the region shown. Black lines show locations of telomere contacts at 25 kb resolution. **(B)** SPRITE bins containing centromere sequence have higher telomere contact frequencies than those that do not contain centromere sequence (unpaired t-test, p<0.001 for all comparisons). Bins with a contact frequency of exactly zero were excluded under the assumption that they lack mappable DNA. **(C)** Telomere-centromere FISH in U2OS reveals many proximal telomere and centromere foci (yellow arrows annotate examples). **(D)** Telomeres and centromeres spatially associate more than expected by random chance. Percentile shows what percentile the mean number of adjacent telomeres per centromere in a given nucleus would be compared to a distribution of adjacent telomere rates when randomizing telomere positions. Each dataset consists of more than 100 nuclei.

To validate the telomere-centromere association observed in SPRITE data, we performed telomere and centromere FISH on a panel of several cell lines with a C-rich telomere probe and a universal centromere probe complementary to the CENP-B binding sequence (Figure 3C). To determine if telomeres and centromeres are spatially proximal, we calculated the average number of nearby telomere foci per centromere focus in a given nucleus. We then randomized the positions of the telomeres 1000 times and determined the percentile of proximal telomeres compared to the random distribution. We observed that nuclei generally exhibit higher rates of telomere-centromere association than the 50^th^ percentile of the randomized distribution, suggesting a spatial association between the two (Figure 3D).

### Spatial association of telomeres with heterochromatin

Genomic regions that share similar chromatin states are thought to spatially associate.^14,15,24^ We hypothesized that the spatial association between telomeres and centromeres may be reflective of a broader tendency for telomeres to spatially associate with other heterochromatic regions.

We found the strongest evidence for a general relationship between telomere contacts and heterochromatin in GM12878; in contrast, ALT cell lines did not show consistent evidence of this trend. A/B compartment eigenvectors provide one method of quantifying the chromatin state of genomic loci. By convention, a more positive eigenvector indicates a stronger association with the A (euchromatic) compartment, whereas a more negative eigenvector indicates a stronger association with the B (heterochromatic) compartment. GM12878 exhibited a robust negative correlation between telomere contacts and the compartment eigenvector, indicating enhanced telomere contacts with heterochromatin. In contrast, other cell lines displayed weaker or insignificant correlations (Figure 4A). We also determined if SPRITE contacts within telomere-containing clusters were enriched at close proximity to H3K9me3 peaks as identified by ChIP-seq in GM12878 (ENCODE)^25–27^ or CUT&Tag in U2OS, G292-iDAXX ALT on/off, and HOS. Substantial spikes in telomere contacts near H3K9me3 peaks are seen in U2OS and HOS, with a weak spike in GM12878 and less evidence of a spike in the two G292-iDAXX treatments (Figure 4B).

**Figure 4.**
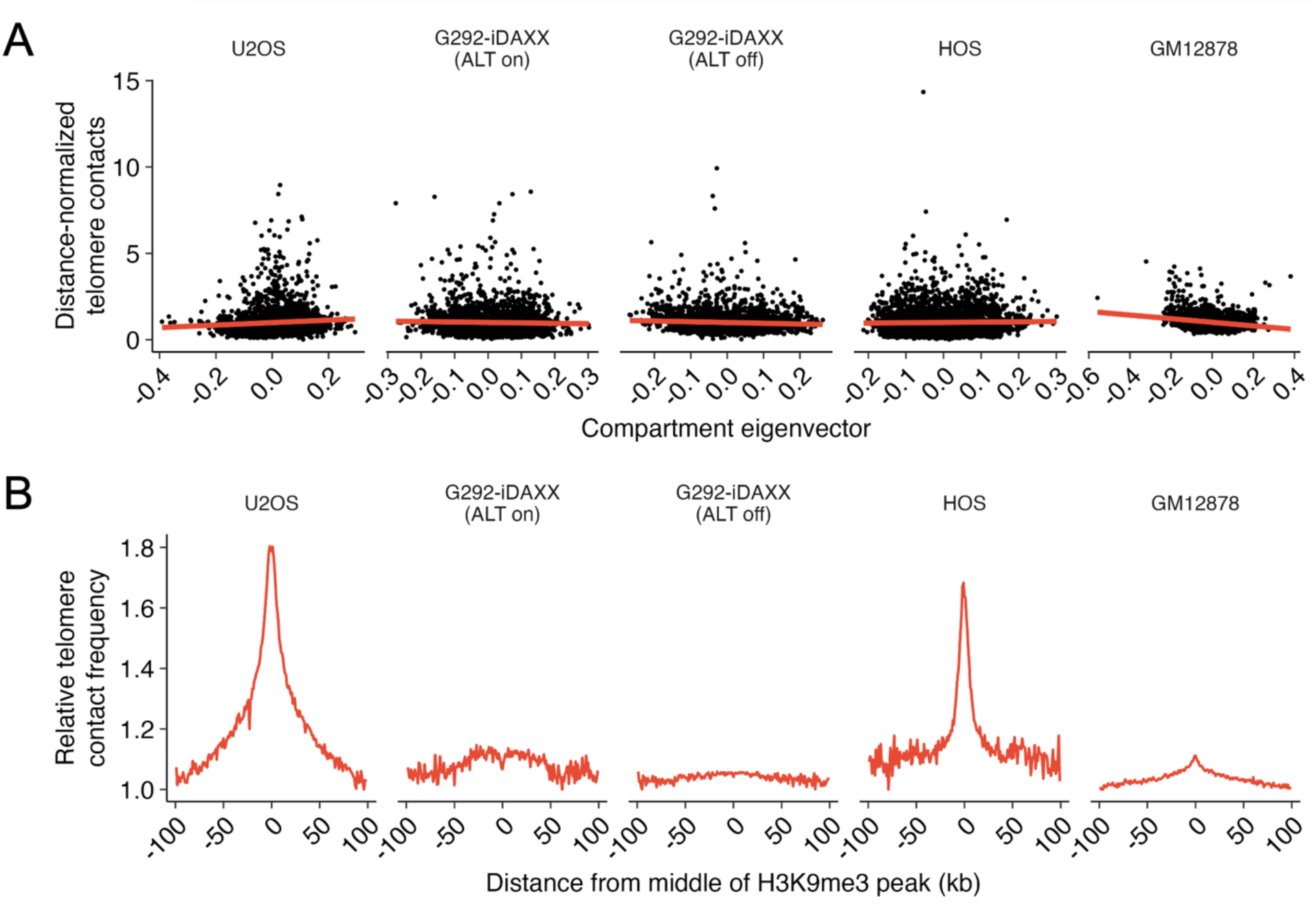
Relationship between telomere contacts and heterochromatin is most evident in GM12878. **(A)** Negative relationship between telomere contacts and compartment eigenvector is robust for GM12878. For G292-iDAXX (ALT off), the relationship is weak and negative whereas it is weak and positive for U2OS. G292-iDAXX (ALT on) and HOS do not show significant relationships. **(B)** Telomere contact frequencies compared to distance from the center of H3K9me3 peaks identified by ChIP-seq for GM12878 (ENCODE) and CUT&Tag for other cell lines. U2OS and HOS display robust spikes in telomere contacts at H3K9me3 peaks, GM12878 shows a small spike, and both treatments of G292-iDAXX show less clear evidence of a spike. Telomere contact frequencies were determined by calculating the ratio of the number of reads aligning at a given distance from the nearest H3K9me3 peak in SPRITE clusters containing telomeres divided by the number of reads at the same distance in SPRITE clusters without telomeres. The minimum value was normalized to one.

### Structural variants containing telomere repeats in ALT cell lines

ALT-positive tumors often have structural variants containing telomere repeats.^13^ A recent analysis of whole genome long read sequencing data from U2OS revealed the presence of dozens of such variants.^28^ In our SPRITE data, we observed a large number of “spikes” in telomere contact frequency at loci internal to the chromosome in U2OS and G292-iDAXX (Figure 2A). As previously stated, loci linearly proximal to telomere repeats are expected to exhibit enhanced contacts in 3-dimensional space with telomere repeat fragments as measured by TelSPRITE. Furthermore, since ALT cell lines have many ectopic telomere sequences, we hypothesized that these loci with enhanced telomere contact frequencies may represent structural variants containing telomere repeats. To test this hypothesis, we performed long read sequencing and utilized a customized pipeline to search for structural variants that include telomere sequence.

Briefly, we searched for reads that contained at least one 300 bp or longer region of soft-clipped sequence that started at a canonical telomere repeat and contained at least 45 telomere repeats total (variant or canonical). Reads with low mapping quality scores, alignments very close to the chromosome end, and alignments in poorly mapped regions were removed from consideration (see Methods). Loci that had at least two reads satisfying the aforementioned criteria were considered to have ectopic telomere repeats. This method serves as a general search for structural variants containing telomere repeats in soft-clipped sequence. These variants could appear as “neotelomeres” with long tracts of telomere repeats that terminate abruptly or may be more complex rearrangements in which other sequences are mixed in with telomere repeats.

We determined that 80 loci in U2OS and 50 loci in G292-iDAXX met our standards to count as sites with ectopic telomere repeats. As a point of comparison with a non-ALT cell line, we analyzed an ONT dataset from the fibroblast cell line HS-5 and only identified three events that met our standards to count as ectopic telomere repeats (Figure 5A, Table S2). SPRITE bins that overlap ectopic telomere repeats in U2OS and G292-iDAXX had markedly higher telomere contact frequencies than those that did not (Figure 5B). Examining individual ectopic repeat locations in both U2OS and G292-iDAXX showed enhanced telomere contact frequencies at close linear distance to the ectopic repeats, underscoring the ability of TelSPRITE to accurately determine which genomic loci are linearly proximal to telomere repeats (Figure 5C-D). Ectopic telomere repeats generally appeared as large tracts of mostly canonical telomere repeats that extended for up to a few kilobases before ending abruptly (Figure S3A,B).

**Figure 5.**
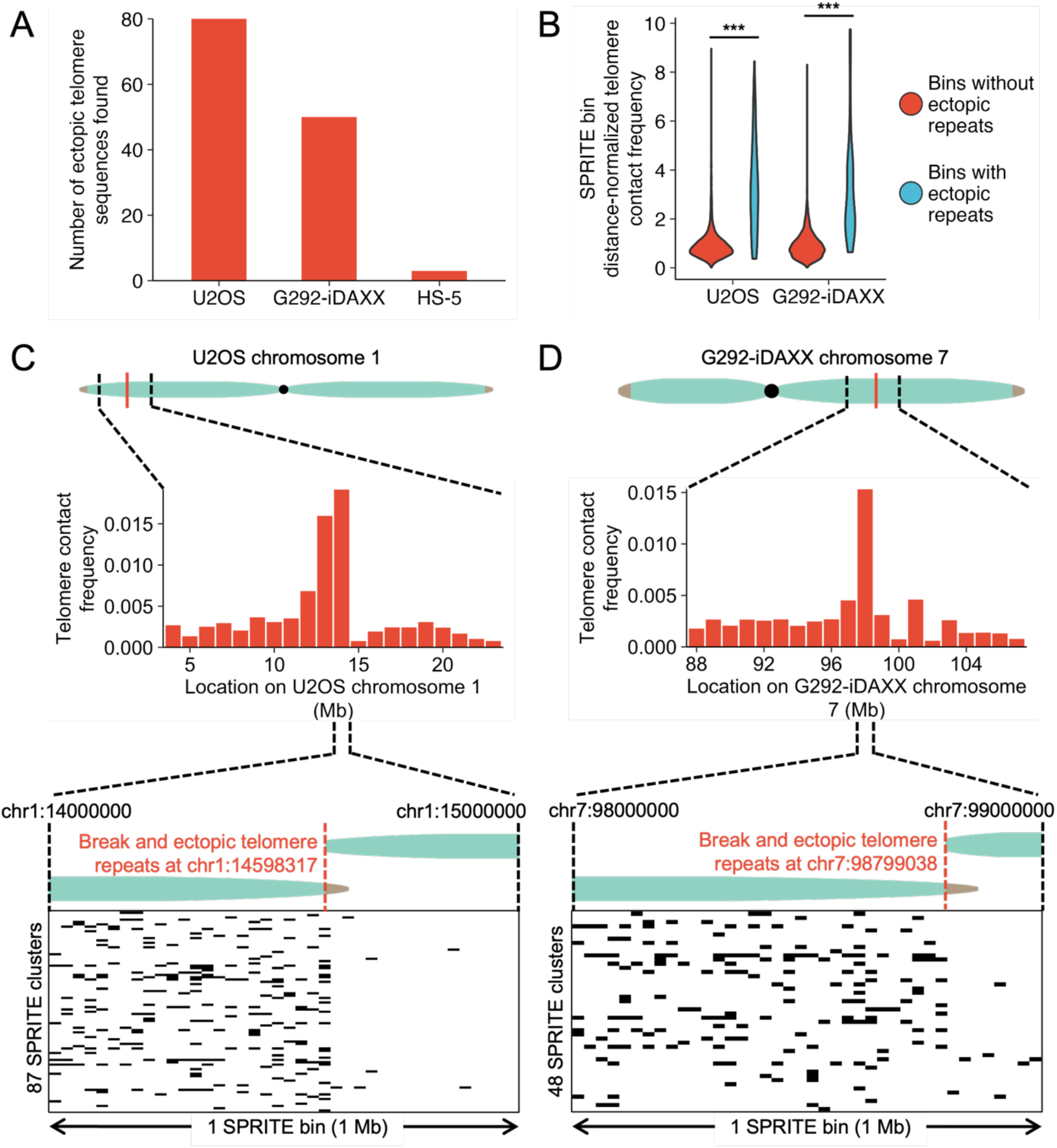
Spikes in telomere contacts reveal ectopic telomere repeats in ALT cell lines. **(A)** Number of ectopic telomere repeats detected in U2OS, G292-iDAXX, and HS-5 by filtering ONT long read sequencing data for reads containing soft-clipped telomere sequence. **(B)** SPRITE bins containing ectopic telomere repeats are substantially enriched for telomere contacts compared to those not containing ectopic telomere repeats in both U2OS and G292-iDAXX (unpaired t-test, p<0.001 for both comparisons). **(C)** Zooming in on the area surrounding the ectopic telomere repeat at chr1:14598317 in U2OS shows high a telomere contact frequency around the addition site (top). Cluster plot at 25 kb resolution of individual SPRITE clusters of size 100 or lower with telomere contacts in the bin containing the ectopic telomere repeat show frequent contacts near the ectopic telomere repeat (bottom). **(D)** Telomere contacts are high in G292-iDAXX near an ectopic telomere repeat at chr7:98799038 (top). Cluster plot at 25 kb resolution of SPRITE clusters size 1000 or lower shows large amounts of telomere contacts near the ectopic telomere repeat site (bottom).

We had hypothesized that ectopic telomere repeats result from recombination events with the telomeres and that genomic loci that are particularly likely to be proximal to the telomeres in 3-dimensional space would be more likely to acquire ectopic telomere repeats. Loci most likely to spatially contact telomeres include those at close linear proximity to chromosome ends and centromeres, so our hypothesis would predict that ectopic telomere repeats would be disproportionately close on the chromosome to these loci. Comparing the locations of ectopic telomere repeats to randomly selected loci showed that the additions were slightly closer to chromosome ends and centromeres than expected by random chance in U2OS, but the same association was not found in G292-iDAXX (Figure S3C,D).

### Rapid ALT suppression causes few rearrangements in telomere contacts

We originally hypothesized that rapid suppression of ALT would result in substantial remodeling of telomere contacts. We found few substantial changes in telomere contact frequencies with suppression of ALT in G292-iDAXX. Examining specific loci revealed that telomere contact frequencies at the 1 Mb bin level typically were relatively consistent regardless of ALT status (Figure 6 top). Additionally, examining telomere contacts near the read level revealed that telomere contacts occur in similar locations (Figure 6 bottom).

**Figure 6.**
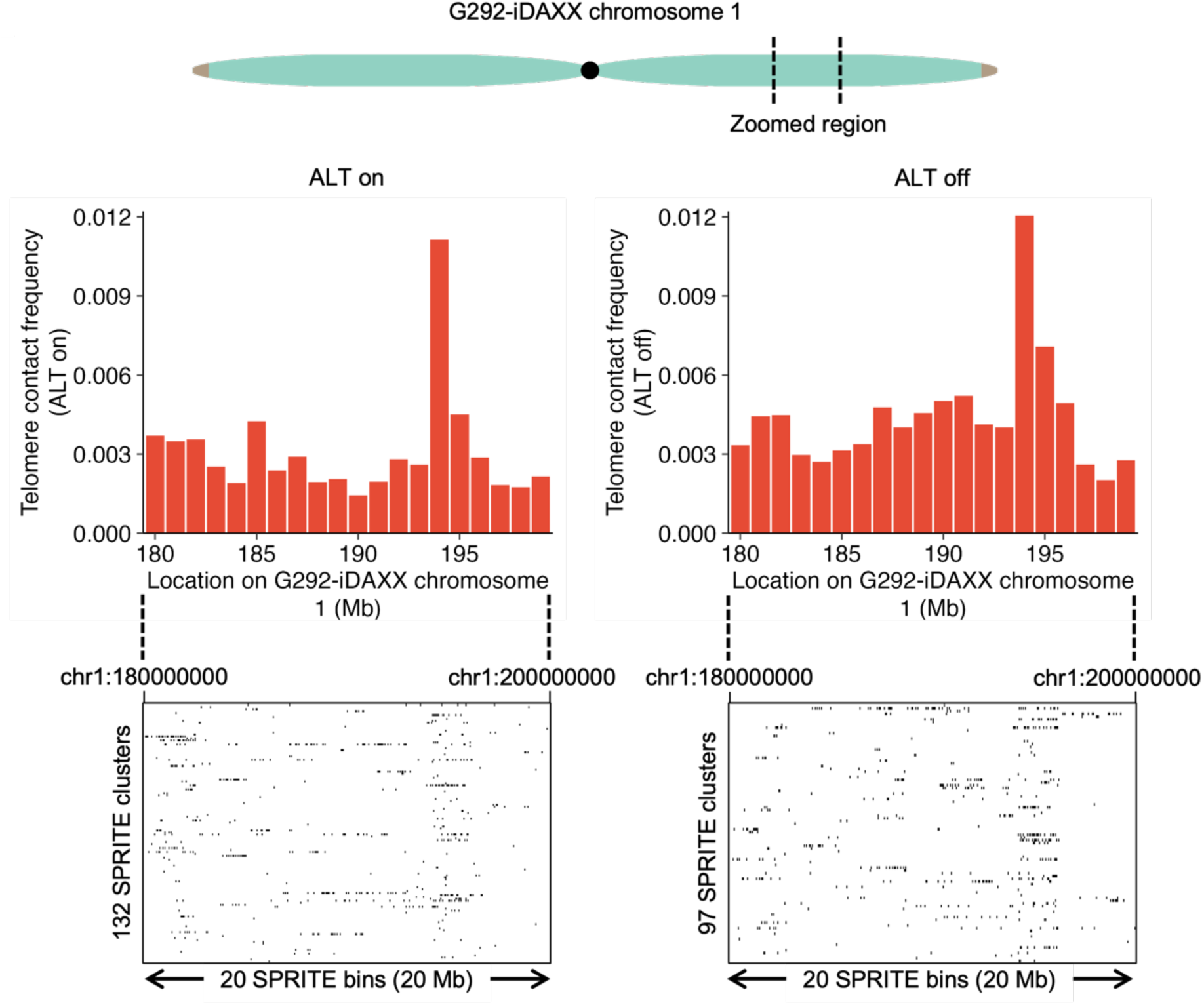
ALT suppression causes minimal change in telomere contacts in G292-iDAXX. Examination of a region of the q-arm of chromosome 1 shows that telomere contacts are broadly similar with and without ALT active. Bar chart shows telomere contact frequencies at 1 Mb resolution and cluster plot shows telomere contacts in SPRITE clusters of size 100 or lower at a resolution of 50 kb.

However, rapid suppression of ALT did lead to a noticeable decline in the percentage of reads passing the telomere filter (Figure 1D). Measurable telomere length attrition is not expected over the short time period used for ALT suppression nor did we observe any (Figure 1C). Therefore, the decline in telomere read counts may be explained by loss of ALT-specific extrachromosomal telomere DNA.

## Discussion

Genomic rearrangement events involving telomeres represent a significant source of genomic instability in ALT tumors; however, little is known about what genomic loci are prone to undergo these rearrangements and how they occur. We hypothesized that genomic loci which frequently contact telomeres in space are likely to experience recombination events with telomeres. However, existing computational workflows for 3-dimensional genomics assays do not enable assessment of the spatial distribution of telomeres, as repetitive regions are typically ignored. In order to quantify telomere contact frequencies, we developed TelSPRITE, a modified version of the SPRITE analysis pipeline that extracts telomere contacts.

We found a consistent spatial association between telomeric and centromeric chromatin regardless of telomere maintenance type. This association was the most visually prominent feature in the telomere contact pattern besides the linear dependance of telomere contact frequency on the distance from the chromosome end. Regions with similar chromatin states tend to spatially associate,^14,15,24^ so this relationship may reflect spatial clustering of telomeric and centromeric constitutive heterochromatin. Interestingly, the wider relationship between telomere contact frequency and heterochromatin across the genome was murkier, especially in the ALT cell lines, with correlations between telomere frequency and markers of heterochromatin often being weak or zero. ALT telomeres are known to have a substantially altered chromatin state characterized by telomere decompaction,^29,30^ so this diminished relationship may result from disruption of telomeric heterochromatin.

We wished to determine which parts of the genome were most likely to undergo recombination with telomeres in ALT tumors. Initially, we had anticipated that we would find that telomeres were spatially rearranged in ALT. Furthermore, we thought that loci that were spatially proximal to telomeres in ALT would be likely to experience recombination events with them, given that ALT telomeres continuously undergo a process of homologous recombination. We found few noteworthy trends in telomere rearrangements with ALT but instead found that ectopic telomere repeats themselves had a particularly large impact on the telomere contact pattern in ALT cells. The number of these ectopic telomere repeats and their impact on the TelSPRITE contact pattern was larger than anticipated. A recent study searched for ectopic telomere repeats in multiple cell lines, including U2OS, and uncovered large numbers of neotelomeres and a smaller number of chromosomal fusion events.^28^ Our results are in agreement that ALT cell lines have many structural variants containing telomere repeats. Furthermore, our TelSPRITE approach provides an alternative method of identifying candidate sites of ectopic telomere repeats that can augment long read sequencing data.

Unexpectedly, rapid suppression of ALT was associated with a substantial decline in telomere read counts. The existence of extrachromosomal DNA species, particularly C-circles, is a hallmark of ALT.^31–33^ C-circles largely disappear with a few days of ALT suppression in G292-iDAXX;^18^ therefore, a decline in bulk telomere content may reflect loss of this extrachromosomal telomere DNA. Alternatively, changes in the chromatin state at the telomeres could also affect the number of DNA fragments from telomeres in the SPRITE library. Open chromatin is generally understood to yield more DNA fragments in chromatin profiling assays, including Hi-C, since it is more efficiently fragmented.^34,35^ Compaction of telomeres with ALT suppression may therefore decrease sequencing coverage of telomeres in SPRITE. Notably, this hypothesis is not mutually exclusive with the hypothesis that loss of C-circles accounts for some of the decline in telomere reads.

TelSPRITE enables determination of telomere contact frequencies with simple modifications to the computational workflow of SPRITE. Our method comes with both benefits and limitations. Since we did not modify the assay, which yields genome-wide chromatin interactions, we were able to extract telomere contact frequencies from the same dataset that we used to assess other principles of chromatin organization. This was particularly useful for certain downstream analysis applications, such as the comparisons between telomere contact frequencies and A/B compartmentalization provided in this work. Furthermore, our computational pipeline could easily be adapted to profile spatial contacts of other repeat regions by simply changing the filtering script to search for reads consisting of a different repeat. The primary drawback with TelSPRITE is sequencing inefficiency if the primary goal is to only assess telomere contacts, as only a small subset of SPRITE clusters contain telomeres. SPRITE has previously been adapted to include enrichment procedures for clusters containing proteins of interest.^36^ An enrichment procedure for telomeres, such as a pulldown of SPRITE clusters containing a telomere binding protein, may help improve telomere counts and resolution. Quantifying telomere contact frequencies at enhanced resolution may enable detection of ALT-specific rearrangements that were not visible in the present study.

## Methods

### Cell culture

U2OS and G292-iDAXX cell lines were maintained in McCoy’s 5A modified media with 15% fetal bovine serum. HOS, MG-63, and hTERT RPE-1 were grown in RPMI, Eagle’s Minimum Essential Media, and DMEM-F12 respectively, all with 10% fetal bovine serum. Penicillin/streptomycin (1%) and/or Gibco antibiotic-antimycotic (1X or 2X concentration) were added to all media.

G292-iDAXX was maintained in 100 μg/mL Gibco Geneticin. Suppression of ALT in G292-iDAXX was performed by inducing expression of wild-type DAXX with 10 ng/mL doxycycline for five days. Successful suppression of ALT was confirmed via a C-circle assay.

### Southern blot

Agarose plugs for pulse field agarose gel were prepared using the BioRad CHEF Mammalian Genomic DNA Plug Kit. Adherent cells were trypsinized and counted, and 1% agarose plugs were prepared with 5 million cells each. Plugs were digested with proteinase K at 50°C, washed in wash buffer supplemented with 1 mM PMSF, and digested in HinfI and RsaI at 37°C overnight prior to running the gel. DNA was separated via pulse-field electrophoresis, using a 1% agarose gel, in 0.5X TBE. The gel was run at 6 V cm^-1^ for 15 h with switch times of 0.1-6.0 s. After separation, the gel was blotted to Nytran SPC charged membrane (Whatman) and was probed with telomere specific 3’ labeled DIG probes (CCCTAA)_3_ (IDT) used at a concentration of 15 ng/mL. Detection was accomplished using the DIG DNA Labeling and Detection Kit (Sigma-Aldrich).

### SPRITE assay

We performed the SPRITE assay according to the published protocol.^17^ Adherent cells were trypsinized and then crosslinked in suspension. We utilized the suggested two-step crosslinking procedure in which cells are first crosslinked with 2 mM DSG for 45 minutes before 10 minutes of crosslinking in 3% formaldehyde. We divided our crosslinked pellets into aliquots of 10 million cells each and performed the remainder of the assay with one of these aliquots.

Sonication conditions for SPRITE must be optimized to produce a range of cluster sizes and DNA fragments of a length appropriate for sequencing. For U2OS and G292-iDAXX experiments, cell pellets were sonicated on a Covaris S2 sonicator for 11 or 12 cycles with the following parameters: 10% duty cycle, intensity of 5, and 200 cycles/burst. Cycles lasted 30 seconds each with a 10 second rest between them. The HOS experiment utilized a Covaris e220 sonicator; duty factor was set to 12% and peak power (which replaces the intensity parameter) was set to 145 W while all other settings remained the same. Following sonication, a DNase titration procedure was utilized to digest DNA fragments down to a size optimal for Illumina next-generation sequencing (about 100 to 1000 base pairs). We utilized a coupling ratio (the ratio of the number of DNA fragments to the number of NHS beads) of 1 or lower for all experiments to minimize false positive interactions resulting from SPRITE clusters coupling to the same beads.

SPRITE libraries were sequenced on an Illumina NextSeq 2000 with at least 500 million raw sequencing reads per data set. To ensure that we captured most reads within each SPRITE cluster, we estimated the number of unique DNA molecules in our libraries based on library molarity and the number of PCR cycles used in the amplification and sequenced at >1X coverage.

### Extracting telomere contacts from SPRITE data

Telomere contact frequencies were determined from SPRITE data using a modified version of the published SPRITE analysis pipeline.^17^ SPRITE utilizes iterative barcoding to tag each DNA molecule in a cluster of crosslinked genomic DNA fragments with a “barcode” that is specific to that cluster. Paired end sequencing captures both the genomic DNA fragment and the barcode, so interactions can be inferred by aligning DNA fragments to a reference genome and then determining which fragments share barcodes. Contact frequency is mostly a function of the rate at which reads aligning to any two genomic loci can be found together in the same SPRITE clusters. We retained the “2/N” downweighting procedure, meaning that the effect of an interaction within a cluster on the contact frequency is inversely proportional to the size of the cluster. A 2-dimensional matrix is generated in which each row represents a genomic locus (or “bin”) consisting of a segment of a chromosome of a given size. We used a resolution of 1 Mb for our analyses, so each row in the contact matrix represents a 1 Mb segment of a chromosome. Each position in the matrix contains the contact frequency between the two genomic loci represented by the rows intersecting at that position.

Our approach to quantify telomere contact frequencies was to treat telomeres as their own “locus” and include that locus during generation of the contact matrix. After some initial preprocessing steps such as trimming of adapter sequences, we utilized a custom Python script that sorted through the sequences of reads in fastq files to determine if they had enough telomere repeats to count as a telomere. Unless otherwise indicated, all telomere contact data we show were filtered at “medium stringency” conditions, meaning that reads counted as a telomere if they had at least 4 canonical telomere repeats (TTAGGG/CCCTAA) and at least 7 total telomere repeats when variant repeats were included. If a read was determined to be a telomere, it was added to a bam file and its location was manually assigned to the chrT locus. Reads determined to not be telomeres were aligned to the hg38 reference genome as normal and added to a separate bam file. These bam files were subsequently recombined into a single bam file that underwent masking of repetitive elements (which does not affect chrT reads) and was used to generate a cluster file containing the read positions in each SPRITE cluster. The cluster file was used to generate the contact frequency matrix, and then the telomere contact frequencies stored in the row of the matrix corresponding to chrT could be extracted and utilized for subsequent analyses.

### Determining A/B compartment eigenvectors from SPRITE contact matrices

A/B compartments are usually assessed via a principal component analysis (PCA) of SPRITE/Hi-C contact matrices.^37^ In our experience, looking at the first principal component of the PCA was usually not sufficient to generate a compartment signal with positive/negative oscillations that matched the “plaid” pattern in SPRITE matrices. Typically, the first principal component would label the chromosome arms or some other irrelevant feature, and examining the second, third, fourth, or fifth principal component was necessary.

We sought to apply consistent criteria for which principal component was selected to represent the compartment signal in SPRITE matrices. First, we converted the raw SPRITE matrices into Pearson correlation matrices like those shown in Figure S1C. We used the same process previously described for Hi-C matrices in which each contact frequency is first divided by the genome-wide average contact frequency between loci at the same linear distance.^14^ Next, we calculated the Pearson correlation between the rows that intersected at a given position in the matrix. We excluded loci that mostly had contact frequencies of zero when computing the correlation.

Finally, we subjected the matrices to a PCA. For each of the first five principal components, we examined the gene densities of the two compartments consisting of loci with the same sign (positive or negative). The first component that yielded compartments whereby one compartment had 1.5 times the gene density of the other was selected to represent the A/B compartment eigenvector. If none of the first five principal components met the gene density cutoff, then the one with the largest discrepancy in gene density was selected. Signs were reversed if necessary to ensure that the more gene-dense compartment (A compartment) had a positive sign. This procedure was used because preliminary testing showed it to yield eigenvectors that visually matched the plaid pattern seen in the Pearson correlation matrices.

### Determining locations of ectopic telomere repeats with long read sequencing

DNA for Oxford Nanopore sequencing was prepared using the Nanobind CBB Kit (PacBio) according to the manufacturer’s protocol. The single G292-iDAXX library, the single HS-5 library, and one of two U2OS libraries were prepared by first treating 3 μg of DNA with the Short Fragment Eliminator Kit (ONT) according to the manufacturer’s protocol before shearing with the DNA Fluid+ Kit on a Megaruptor 3 (Diagenode) to a mean size of approximately 50k bp. The libraries were prepared using the Ligation Sequencing Kit V14 (ONT), according to the manufacturers protocol. A second U2OS library was prepared from 30 μg of DNA using the Ultra-Long Sequencing Kit (ONT) according to the manufacturer’s protocol. Libraries were sequenced using a PromethION 2 Solo sequencer on a R10.4.1 Flow Cell.

Base calling was performed using Dorado and reads were subsequently aligned to the hg38 reference genome using minimap2.^38,39^ The aligned bam files from the two U2OS libraries were combined for downstream analysis.

After aligning long reads to the reference genome, locations of interstitial telomere sequence not found in the reference genome were extracted with a computational pipeline consisting of a series of custom Python scripts. Reads passed an initial filter for telomere sequence if they contained at least one 300 base pair window starting at a canonical repeat (TTAGGG/CCCTAA) and containing 45 or more total repeats (canonical or variant). Candidate reads passing this filter were subsequently reexamined to determine if the telomere sequence was located in a soft-clipped section of the read; reads in which the telomere sequence was not soft-clipped were assumed to have telomere sequence that is present in the reference genome and were ignored. Reads were classified according to the side of the read that was clipped as well as whether or not the orientation of the telomere repeat sequence was standard or inverted. The insertion location of a telomere repeat sequence was determined based on the start or end position of the read alignment depending on which side of the read had telomere sequence.

Imposition of several conditions was required to ensure that only high confidence ectopic telomere repeats were considered. Reads that passed the telomere filter but had MAPQ scores less than 30 or had telomere insertion locations within 100 kb of the chromosome end were ignored. Additionally, if a telomere insertion locus was poorly mapped (defined as at least 40% of reads overlapping that locus in the original, unfiltered BAM file having a MAPQ score of less than 10), that locus was also discounted.

Finally, we limited our analysis to loci that were supported by at least two reads. To account for the fact that sometimes two reads may be at nearly the same locus but have their alignment start or end point differ by a few base pairs, reads whose start positions differed by 100 or fewer base pairs were considered to correspond to the same locus.

Ectopic telomere repeats were considered to intersect a gene if the start position fell anywhere within the transcriptional window of that gene.

### Telomere-centromere FISH

Telomere-centromere FISH was performed by growing cells on acid-washed coverslips and then fixing in 4% formaldehyde for 15 minutes. Cells were permeabilized in 0.5% Triton X-100 for 5 minutes, washed in PBS, and treated with 100 μg/mL RNase A at 37°C for 20 to 30 minutes. Cells were washed again with PBS and then dehydrated.

Probes were TelC-Cy3 and CENPB-FAM, both from PNA Bio. Probes were added together (each at a 1:50 dilution) to hybridization buffer (70% deionized formamide, 10 mM Tris-HCl pH 7.5, 0.5% 10x Roche blocking solution). Diluted probes were denatured at 90°C and coverslips were denatured at 85°C, both for 5 minutes. Coverslips were subsequently flipped onto an aliquot of diluted probe, and the probes and coverslips were denatured together at 85°C for 5 minutes. Coverslips were incubated on the probes at room temperature overnight in a humidified chamber. Coverslips were subsequently washed in formamide-Tris solution (70% deionized formamide, 10 mM Tris-HCl pH 7.5), washed in TNT (0.05 M Tris pH 7.5, 0.15 M NaCl, 0.05% Tween-20 pH 7.5), and mounted overnight at room temperature on ProLong Glass Antifade Mountant with NucBlue (Thermo Fisher).

### Microscopy

Telomere-centromere FISH images were captured on a Nikon SoRa Spinning Disk microscope with a 60X oil immersion objective. Excitation wavelengths were 405 nm, 488 nm, and 561 nm for DAPI stain, TelC-Cy3 probe, and CENPB-FAM probe respectively. Images in Fig. 3C were cropped, pseudo-colored, smoothed, and adjusted to optimize contrast and brightness using Fiji.

### FISH image analysis

For analysis of telomere-centromere proximity in FISH images, fields of nuclei were first segmented based on DAPI intensity. Threshold intensities were determined for individual image channels using a Multi-Otsu algorithm, and foci with a minimum of nine pixels in area were segmented. Telomere foci within five pixels of a centromere focus were scored as centromere-adjacent. To test the hypothesis that telomeres were disproportionately located adjacent to centromeres, a randomization approach was utilized. For each nucleus image, telomere foci were programmatically randomized within the nuclear area 1000 times, and telomere-centromere proximity was scored for each randomization. The actual proximity was assigned a percentile rank in comparison with the 1000 randomizations.

### CUT&Tag

CUT&Tag was performed using a commercial kit from EpiCypher and a polyclonal anti-H3K9me3 antibody (Active Motif). This antibody was selected after preliminary CUT&Tag experiments with low sequencing depth revealed it to yield high concentrations of mononucleosome-size fragments. Crucially, it was the most specific to the H3K9me3 target out of four anti-H3K9me3 antibodies initially tested (EpiCypher provides a “K-MetStat Panel” of custom nucleosomes that can be spiked into CUT&Tag samples to test specificity of antibodies against histone post-translational modifications).

For each CUT&Tag reaction, 100,000 cells were used. The protocol provided by EpiCypher was followed without modification. Negative and positive controls with rabbit IgG and anti-H3K27me3 antibodies were performed alongside three technical replicates of the anti-H3K9me3 antibody. Assay success was confirmed by observing enrichment of mononucleosome size fragments in the H3K27me3 and H3K9me3 samples using an Agilent Bioanalyzer or TapeStation. Libraries were sequenced at about 20 million raw reads per library on an Illumina NextSeq 2000.

### Analysis of CUT&Tag data

CUT&Tag reads first underwent a local pairwise alignment to the hg38 reference genome using Bowtie 2. Only reads with a MAPQ score of at least 20 were retained. Peaks were called using SICER2.^40^

## Acknowledgements

We thank the NCI’s Center for Cancer Research at the NIH Intramural Research Program for supporting this work. We are also grateful for helpful feedback from members of the Meltzer lab, as well as Dr. Eros Lazzerini Denchi and other members of Lazzerini Denchi, Stracker, and Weyemi labs at the CCR. For technical input, training, and assistance with assays and equipment, we thank Dr. Michael Kruhlak and Langston Lim of the CCR Confocal Microscopy Core, Dr. Fan Yang, and David Petersen.

## Author Contributions

D.G.W., S.F.C.S., and P.S.M. designed the project and experiments. D.G.W., S.F.C.S., M.A.P., and R.L.W. performed experiments, and D.G.W., S.F.C.S., and P.S.M. analyzed data. D.G.W., S.F.C.S., and P.S.M. wrote and revised the manuscript.

## Data Availability

Raw data will be made available upon publication.

---

**Figure S1.**
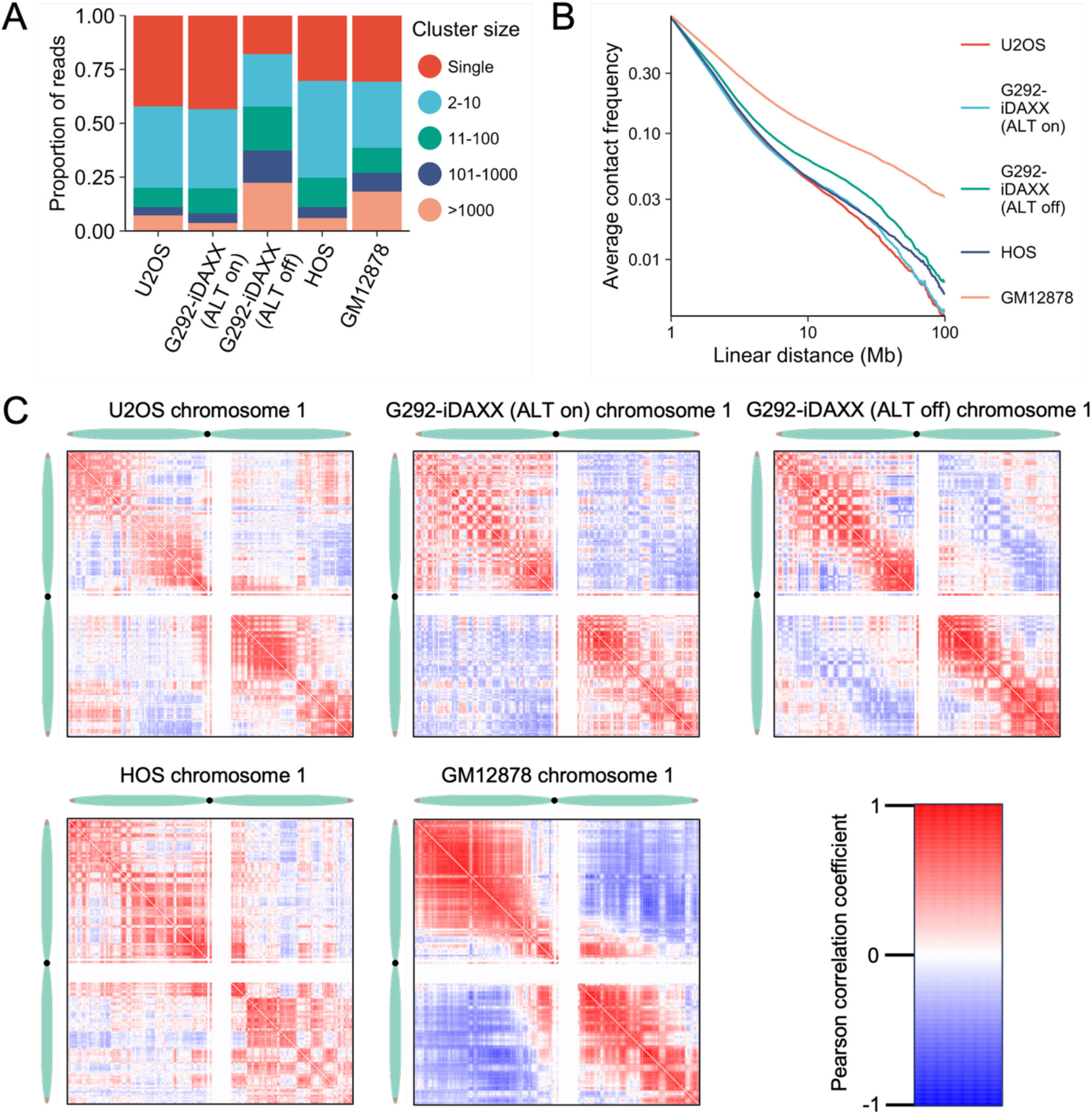
SPRITE captures interactions in clusters of diverse sizes and recapitulates known aspects of genomic spatial organization. **(A)** Cluster size distribution shows that each SPRITE dataset contains reads in both small and large clusters. Bars show the fraction of the total reads in a dataset found in clusters in each size range. **(B)** For all datasets, contact frequency decreases precipitously with increasing chromosomal distance between loci. **(C)** A/B compartments are visible in all datasets. Shown are Pearson correlation matrices of chromosome 1 contacts normalized by linear distance.

**Figure S2.**
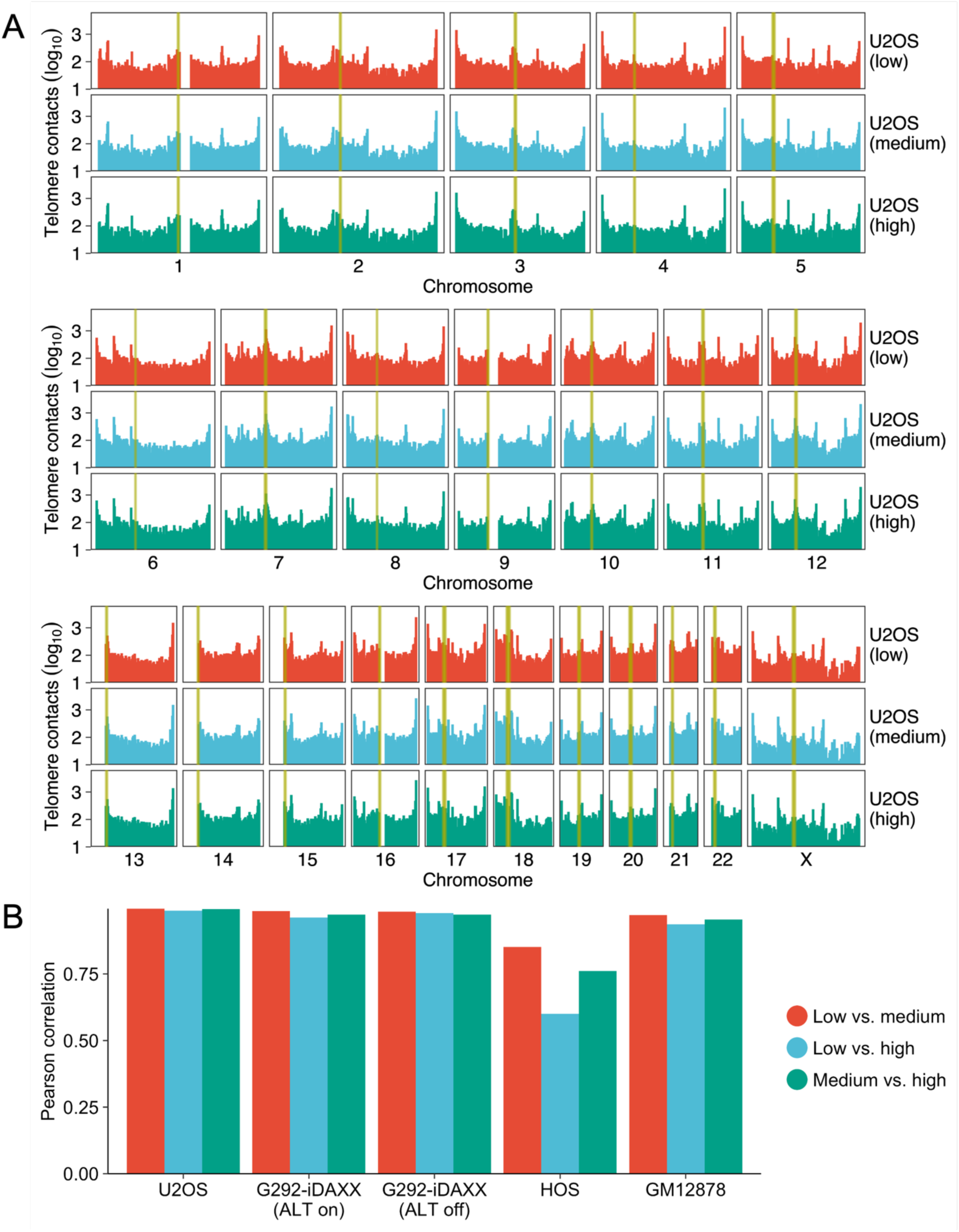
Telomere contact pattern is not substantially influenced by filtering stringency. **(A)** U2OS telomere contacts at three different filtering stringencies: low (4 or more total repeats and 2 or more canonical repeats), medium (7 or more total repeats and 4 or more canonical repeats), and high (14 or more total repeats and 7 or more canonical repeats). **(B)** Telomere contacts calculated at different filtering stringencies correlate strongly. Plot shows the Pearson correlation coefficient between each pair of stringency conditions for each dataset. Bins that had a value of zero in any of the conditions for a given cell line were assumed to lack mappable DNA and were excluded.

**Figure S3.**
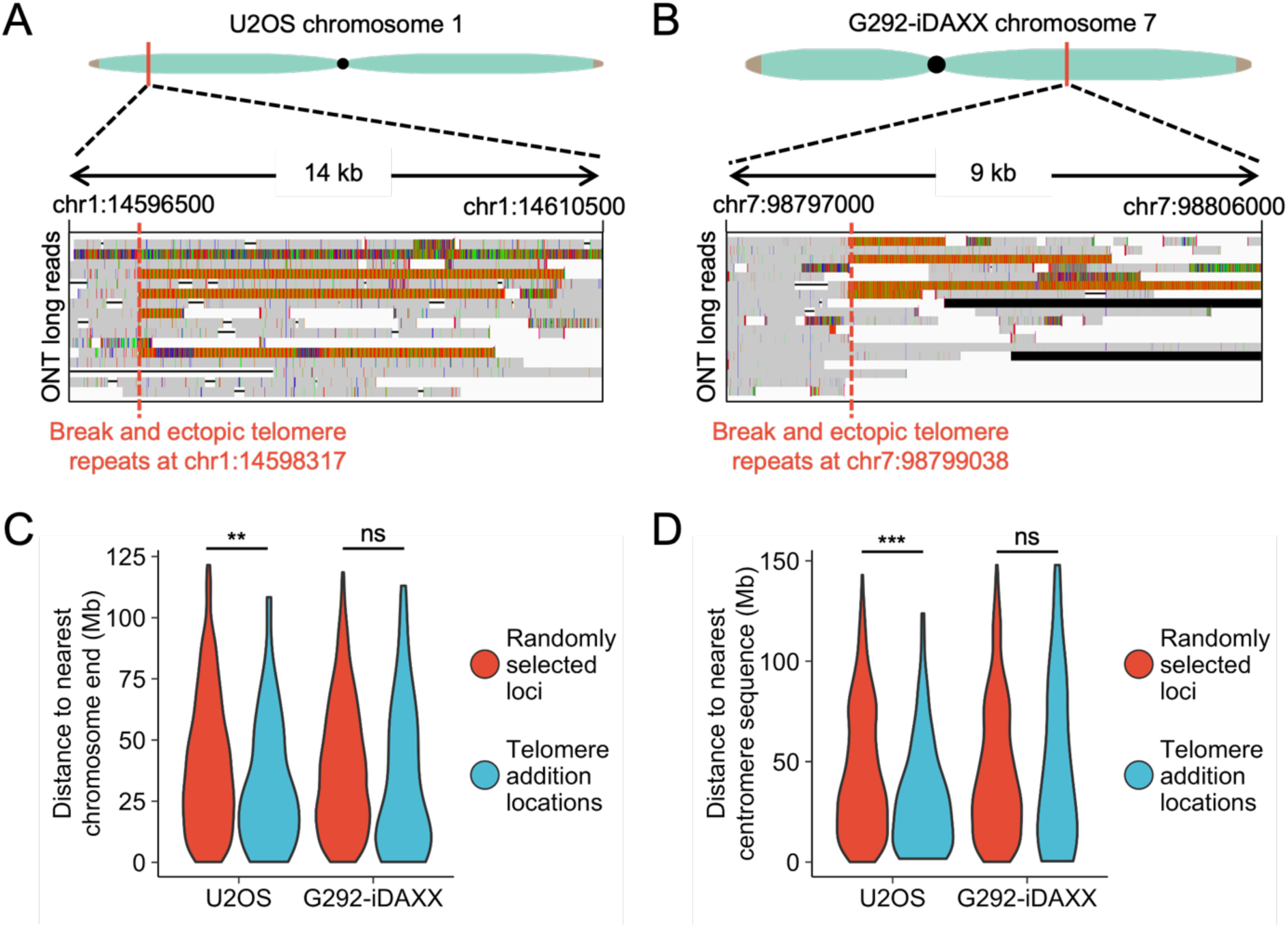
ALT cell lines are characterized by structural variants containing telomere repeats. **(A)** ONT long reads supporting the ectopic telomere repeat in U2OS from Figure 5A. Reads display long segments of telomere repeats that end abruptly, consistent with a neotelomere addition. **(B)** ONT long reads supporting the ectopic telomere repeat in G292-iDAXX from Figure 5B. This structural variant also resembles a neotelomere. **(C)** Ectopic telomere repeats are closer to chromosome ends than expected by random chance in U2OS (unpaired t-test, p<0.01) but not in G292-iDAXX. Loci corresponding to sites of ectopic telomere repeats are compared to set of random loci. For each comparison, 1000 random loci were picked, but some were excluded based on the same criteria used in calling the ectopic telomere repeats so that the exclusion criteria would not bias the analysis. **(D)** Ectopic telomere repeats are closer to centromere sequence than expected by random chance in U2OS (unpaired t-test, p<0.001) but not in G292-iDAXX. Random loci were generated with the same procedure as in (C).

## References

1. Gao J, P.H. (2022). Targeting telomeres: advances in telomere maintenance mechanism-specific cancer therapies. Nat Rev Cancer 22, 515–532. doi: 10.1038/s41568-022-00490-1.

2. Bryan TM, E.A., Gupta J, Bacchetti S, Reddel RR (1995). Telomere elongation in immortal human cells without detectable telomerase activity. EMBO J 14, 4240–4248. doi: 10.1002/j.1460-2075.1995.tb00098.x.

3. Bryan TM, E.A., Dalla-Pozza L, Dunham MA, Reddel RR (1997). Evidence for an alternative mechanism for maintaining telomere length in human tumors and tumor-derived cell lines. Nat Med 3, 1271–1274. doi: 10.1038/nm1197-1271.

4. Dunham MA, N.A., Fasching CL, Reddel RR (2000). Telomere maintenance by recombination in human cells. Nat Genet 26, 447–450. doi: 10.1038/82586.

5. Bower K, N.C., Cole SL, Dagg RA, Lau LM, Duncan EL, Moy EL, Reddel RR. (2012). Loss of wild-type ATRX expression in somatic cell hybrids segregates with activation of Alternative Lengthening of Telomeres. PLoS One 7, e50062. doi: 10.1371/journal.pone.0050062.

6. Heaphy CM, d.W.R., Jiao Y, Klein AP, Edil BH, Shi C, Bettegowda C, Rodriguez FJ, Eberhart CG, Hebbar S, Offerhaus GJ, McLendon R, Rasheed BA, He Y, Yan H, Bigner DD, Oba-Shinjo SM, Marie SK, Riggins GJ, Kinzler KW, Vogelstein B, Hruban RH, Maitra A, Papadopoulos N, Meeker AK (2011). Altered telomeres in tumors with ATRX and DAXX mutations. Science 333, 425. doi: 10.1126/science.1207313.

7. Lovejoy CA, L.W., Reisenweber S, Thongthip S, Bruno J, de Lange T, De S, Petrini JH, Sung PA, Jasin M, Rosenbluh J, Zwang Y, Weir BA, Hatton C, Ivanova E, Macconaill L, Hanna M, Hahn WC, Lue NF, Reddel RR, Jiao Y, Kinzler K, Vogelstein B, Papadopoulos N, Meeker AK; ALT Starr Cancer Consortium (2012). Loss of ATRX, genome instability, and an altered DNA damage response are hallmarks of the alternative lengthening of telomeres pathway. PLoS Genet 8, e1002772. doi: 10.1371/journal.pgen.1002772.

8. Clynes D, J.C., Xella B, Ayyub H, Scott C, Mitson M, Taylor S, Higgs DR, Gibbons RJ (2015). Suppression of the alternative lengthening of telomere pathway by the chromatin remodelling factor ATRX. Nat Commun 6, 7538. doi: 10.1038/ncomms8538.

9. 9. Law MJ, L.K., Voon HP, Hughes JR, Garrick D, Viprakasit V, Mitson M, De Gobbi M, Marra M, Morris A, Abbott A, Wilder SP, Taylor S, Santos GM, Cross J, Ayyub H, Jones S, Ragoussis J, Rhodes D, Dunham I, Higgs DR, Gibbons RJ (2010). ATR-X syndrome protein targets tandem repeats and influences allele-specific expression in a size-dependent manner. Cell 143, 367–378. doi: 10.1016/j.cell.2010.09.023.

10. Clatterbuck Soper SF, M.P. (2023). ATRX/DAXX: Guarding the Genome against the Hazards of ALT. Genes (Basel) 14, 790. doi: 10.3390/genes14040790.

11. Dilley RL, V.P., Cho NW, Winters HD, Wondisford AR, Greenberg RA (2016). Break-induced telomere synthesis underlies alternative telomere maintenance. Nature 539, 54–58. doi: 10.1038/nature20099.

12. Min J, W.W., Shay JW (2017). Alternative Lengthening of Telomeres Mediated by Mitotic DNA Synthesis Engages Break-Induced Replication Processes. Mol Cell Biol 37, e00226–00217. doi: 10.1128/MCB.00226-17.

13. Sieverling L, H.C., Koser SD, Ginsbach P, Kleinheinz K, Hutter B, Braun DM, Cortés-Ciriano I, Xi R, Kabbe R, Park PJ, Eils R, Schlesner M; PCAWG-Structural Variation Working Group; Brors B, Rippe K, Jones DTW, Feuerbach L; PCAWG Consortium (2020). Genomic footprints of activated telomere maintenance mechanisms in cancer. Nat Commun 11, 733. doi: 10.1038/s41467-019-13824-9.

14. Lieberman-Aiden E, v.B.N., Williams L, Imakaev M, Ragoczy T, Telling A, Amit I, Lajoie BR, Sabo PJ, Dorschner MO, Sandstrom R, Bernstein B, Bender MA, Groudine M, Gnirke A, Stamatoyannopoulos J, Mirny LA, Lander ES, Dekker J (2009). Comprehensive mapping of long-range interactions reveals folding principles of the human genome. Science 326, 289–293. doi: 10.1126/science.1181369.

15. Quinodoz SA, O.N., Tabak B, Palla A, Schmidt JM, Detmar E, Lai MM, Shishkin AA, Bhat P, Takei Y, Trinh V, Aznauryan E, Russell P, Cheng C, Jovanovic M, Chow A, Cai L, McDonel P, Garber M, Guttman M (2018). Higher-Order Inter-chromosomal Hubs Shape 3D Genome Organization in the Nucleus. Cell 174, 744–757. doi: 10.1016/j.cell.2018.05.024.

16. 16. Beagrie RA, S.A., Schueler M, Kraemer DC, Chotalia M, Xie SQ, Barbieri M, de Santiago I, Lavitas LM, Branco MR, Fraser J, Dostie J, Game L, Dillon N, Edwards PA, Nicodemi M, Pombo A (2017). Complex multi-enhancer contacts captured by genome architecture mapping. Nature 543, 519–524. doi: 10.1038/nature21411.

17. Quinodoz SA, B.P., Chovanec P, Jachowicz JW, Ollikainen N, Detmar E, Soehalim E, Guttman M (2022). SPRITE: a genome-wide method for mapping higher-order 3D interactions in the nucleus using combinatorial split-and-pool barcoding. Nat Protoc 17, 36–75. doi: 10.1038/s41596-021-00633-y.

18. Yost KE, C.S.S., Walker RL, Pineda MA, Zhu YJ, Ester CD, Showman S, Roschke AV, Waterfall JJ, Meltzer PS (2019). Rapid and reversible suppression of ALT by DAXX in osteosarcoma cells. Sci Rep 9, 4544. doi: 10.1038/s41598-019-41058-8.

19. Conomos D, S.M., Hills M, Neumann AA, Bryan TM, Reddel RR, Pickett HA (2012). Variant repeats are interspersed throughout the telomeres and recruit nuclear receptors in ALT cells. J Cell Biol 199, 893–906. doi: 10.1083/jcb.201207189.

20. Lee M, H.M., Conomos D, Stutz MD, Dagg RA, Lau LM, Reddel RR, Pickett HA (2014). Telomere extension by telomerase and ALT generates variant repeats by mechanistically distinct processes. Nucleic Acids Res 42, 1733–1746. doi: 10.1093/nar/gkt1117.

21. Feuerbach L, S.L., Deeg KI, Ginsbach P, Hutter B, Buchhalter I, Northcott PA, Mughal SS, Chudasama P, Glimm H, Scholl C, Lichter P, Fröhling S, Pfister SM, Jones DTW, Rippe K, Brors B (2019). TelomereHunter - in silico estimation of telomere content and composition from cancer genomes. BMC Bioinformatics 20, 272. doi: 10.1186/s12859-019-2851-0.

22. Nassar LR, B.G., Benet-Pagès A, Casper J, Clawson H, Diekhans M, Fischer C, Gonzalez JN, Hinrichs AS, Lee BT, Lee CM, Muthuraman P, Nguy B, Pereira T, Nejad P, Perez G, Raney BJ, Schmelter D, Speir ML, Wick BD, Zweig AS, Haussler D, Kuhn RM, Haeussler M, Kent WJ (2023). The UCSC Genome Browser database: 2023 update. Nucleic Acids Res 51, D1188–D1195. doi: 10.1093/nar/gkac1072.

23. Miga KH, N.Y., Jain M, Altemose N, Willard HF, Kent WJ (2014). Centromere reference models for human chromosomes X and Y satellite arrays. Genome Res 24, 697–707. doi: 10.1101/gr.159624.113.

24. Falk M, F.Y., Naumova N, Imakaev M, Lajoie BR, Leonhardt H, Joffe B, Dekker J, Fudenberg G, Solovei I, Mirny LA (2019). Heterochromatin drives compartmentalization of inverted and conventional nuclei. Nature 570, 395–399. doi: 10.1038/s41586-019-1275-3.

25. Consortium, E.P. (2012). An integrated encyclopedia of DNA elements in the human genome. Nature 489, 57–74. doi: 10.1038/nature11247.

26. Hitz BC, J.-W.L., Jolanki O, Kagda MS, Graham K, Sud P, Gabdank I, Strattan JS, Sloan CA, Dreszer T, Rowe LD, Podduturi NR, Malladi VS, Chan ET, Davidson JM, Ho M, Miyasato S, Simison M, Tanaka F, Luo Y, Whaling I, Hong EL, Lee BT, Sandstrom R, Rynes E, Nelson J, Nishida A, Ingersoll A, Buckley M, Frerker M, Kim DS, Boley N, Trout D, Dobin A, Rahmanian S, Wyman D, Balderrama-Gutierrez G, Reese F, Durand NC, Dudchenko O, Weisz D, Rao SSP, Blackburn A, Gkountaroulis D, Sadr M, Olshansky M, Eliaz Y, Nguyen D, Bochkov I, Shamim MS, Mahajan R, Aiden E, Gingeras T, Heath S, Hirst M, Kent WJ, Kundaje A, Mortazavi A, Wold B, Cherry JM (2023). The ENCODE Uniform Analysis Pipelines. bioRxiv [Preprint], 2023.2004.2004.535623. doi: 10.1101/2023.04.04.535623.

27. Luo Y, H.B., Gabdank I, Hilton JA, Kagda MS, Lam B, Myers Z, Sud P, Jou J, Lin K, Baymuradov UK, Graham K, Litton C, Miyasato SR, Strattan JS, Jolanki O, Lee JW, Tanaka FY, Adenekan P, O’Neill E, Cherry JM (2020). New developments on the Encyclopedia of DNA Elements (ENCODE) data portal. Nucleic Acids Res 48, D882–D889. doi: 10.1093/nar/gkz1062.

28. Tan KT, S.M., Leibowitz ML, Garrity-Janger M, Shan J, Li H, Meyerson M (2024). Neotelomeres and telomere-spanning chromosomal arm fusions in cancer genomes revealed by long-read sequencing. Cell Genom 4, 100588. doi: 10.1016/j.xgen.2024.100588.

29. Lynskey ML, B.E., Bhargava R, Wondisford AR, Ouriou JB, Freund O, Bowman RW 2nd, Smith BA, Lardo SM, Schamus-Hayes S, Hainer SJ, O’Sullivan RJ (2024). HIRA protects telomeres against R-loop-induced instability in ALT cancer cells. Cell Rep 43, 114964. doi: 10.1016/j.celrep.2024.114964.

30. 30. Episkopou H, D.I., Van Beneden A, Tilman G, Mattiussi M, Gobin M, Arnoult N, Londoño-Vallejo A, Decottignies A (2014). Alternative Lengthening of Telomeres is characterized by reduced compaction of telomeric chromatin. Nucleic Acids Res 42, 4391–4405. doi: 10.1093/nar/gku114.

31. Henson JD, C.Y., Huschtscha LI, Chang AC, Au AY, Pickett HA, Reddel RR (2009). DNA C-circles are specific and quantifiable markers of alternative-lengthening-of-telomeres activity. Nat Biotechnol 27, 1181–1185. doi: 10.1038/nbt.1587.

32. Grudic A, J.-L.A., Haring SJ, Wold MS, Lønning PE, Bjerkvig R, Bøe SO (2007). Replication protein A prevents accumulation of single-stranded telomeric DNA in cells that use alternative lengthening of telomeres. Nucleic Acids Res 35, 7267–7278. doi: 10.1093/nar/gkm738.

33. Nabetani A, I.F. (2009). Unusual telomeric DNAs in human telomerase-negative immortalized cells. Mol Cell Biol 29, 703–713. doi: 10.1128/MCB.00603-08.

34. Imakaev M, F.G., McCord RP, Naumova N, Goloborodko A, Lajoie BR, Dekker J, Mirny LA (2012). Iterative correction of Hi-C data reveals hallmarks of chromosome organization. Nat Methods 9, 999–1003. doi: 10.1038/nmeth.2148.

35. 35. Chandradoss KR, G.P., Kethavath S, Dass M, Singh H, Nayak R, Kurukuti S, Sandhu KS (2020). Biased visibility in Hi-C datasets marks dynamically regulated condensed and decondensed chromatin states genome-wide. BMC Genomics 21, 175. doi: 10.1186/s12864-020-6580-6.

36. Vangala P, M.R., Quinodoz SA, Gellatly K, McDonel P, Guttman M, Garber M (2020). High-Resolution Mapping of Multiway Enhancer-Promoter Interactions Regulating Pathogen Detection. Mol Cell 80, 359–373. doi: 10.1016/j.molcel.2020.09.005.

37. Lajoie BR, D.J., Kaplan N (2015). The Hitchhiker’s guide to Hi-C analysis: practical guidelines. Methods 72, 65–75. doi: 10.1016/j.ymeth.2014.10.031.

38. H, L. (2018). Minimap2: pairwise alignment for nucleotide sequences. Bioinformatics 34, 3094–3100. doi: 10.1093/bioinformatics/bty191.

39. H, L. (2021). New strategies to improve minimap2 alignment accuracy. Bioinformatics 37, 4572–4574. doi: 10.1093/bioinformatics/btab705.

40. Zang C, S.D., Zeng C, Cui K, Zhao K, Peng W (2009). A clustering approach for identification of enriched domains from histone modification ChIP-Seq data. Bioinformatics 25, 1952–1958. doi: 10.1093/bioinformatics/btp340.

